# Maternal Transfer of Oral Vaccine Induced Anti-OspA Antibodies Protects *Peromyscus* spp Pups from Tick-Transmitted *Borrelia burgdorferi*

**DOI:** 10.1101/2025.02.24.639966

**Authors:** Jose F. Azevedo, Greg Joyner, Suman Kundu, Kamalika Samanta, Maria Gomes-Solecki

## Abstract

The efficacy and duration of passive immunity protection depends on maternal antibody levels and transfer efficiency. We investigated whether oral vaccination of *Peromyscus leucopus* dams with recombinant OspA-expressing *E. coli* could induce maternal transfer of anti-OspA antibodies and protect pups from *Borrelia burgdorferi* challenge. Dams were vaccinated until breeding pairs were created (i), until parturition (ii), and until pups were 2 weeks old (iii). Pups were challenged with nymphal *I. scapularis*-transmitted *B. burgdorferi* at ∼ 4 weeks of age. Anti-OspA IgG were quantified in dams and pups, and anti*-B. burgdorferi* IgG were quantified in pups. *B. burgdorferi* burden was assessed by *flaB* qPCR in pups’ tissues ∼ 4 weeks after tick challenge and viability of *B. burgdorferi* was assessed by culture of heart tissue. *P. leucopus* pups born to dams vaccinated until breeding had low serologic anti-OspA antibody and were not protected from tick transmitted *B. burgdorferi* infection. However, when dams’ vaccination extended until parturition and until pups were two weeks old, significant anti-OspA antibody transfer and protection from *B. burgdorferi* infection occurred. This was evidenced by absence of antibody to *B. burgdorferi* PepVF, absence of *B. burgdorferi flaB* DNA in heart and bladder tissues, and absence of *flaB* in culture from heart tissues from pups euthanized >9 weeks after birth. We show that transfer of anti-OspA antibodies from vaccinated *P. leucopus* dams to offspring prevents tick transmission and infection dynamics of *B. burgdorferi* in the major reservoir host of this spirochete in the United States.

**Importance:** This study contributes to our understanding of how interventions based in reservoir targeted OspA vaccines designed to block transmission of *B. burgdorferi* from infected *I. scapularis* ticks may disrupt the enzootic cycle of this spirochete and reduce incidence of Lyme disease.

## Introduction

Lyme disease, caused by *Borrelia burgdorferi*, is the most prevalent vector-borne disease in North America (1). The white-footed mouse (*Peromyscus leucopus*) serves as the primary reservoir host (2), maintaining *B. burgdorferi* in enzootic cycles and facilitating transmission to *Ixodes* ticks that subsequently infect humans and other mammals (3). Unlike humans, reservoir hosts do not develop significant disease following *B. burgdorferi* infection, yet they serve as a crucial source of spirochete acquisition for larval Ixodid ticks (4). Thus, interrupting transmission of *B. burgdorferi* (5) has been proposed as a strategy for controlling Lyme disease (6, 7).

Outer surface protein A (OspA) is a major target for Lyme disease vaccine development (8) due to its essential role in *B. burgdorferi* persistence within the tick vector (9). OspA facilitates spirochete adherence to the tick midgut, allowing the bacterium to survive between blood meals (10). During tick feeding, *B. burgdorferi* downregulates OspA while upregulating OspC (11), which enables migration to the salivary glands and subsequent transmission to the host (12–14). Vaccines targeting OspA aim to elicit antibodies that neutralize spirochetes within the tick gut, preventing their transmission before they reach the mammalian host (15, 16). These vaccines have been explored in multiple contexts, including human vaccination (17, 18), direct vaccination of *P. leucopus* (19), and oral bait delivered reservoir-targeted vaccines (6).

Beyond direct vaccination, maternal transfer of antibody provides an alternative strategy to protect neonates during the critical early stages of life (20). In the context of ecology of vector-borne pathogens, this temporary immunity reduces early-life susceptibility to infection (21). Previous studies have demonstrated that oral vaccination of dams with OspA results in anti-OspA IgG in offspring (22), though the degree to which this protects against *B. burgdorferi* transmission remains unknown. In this study, we addressed this critical gap by investigating the extent of maternal antibody transfer in *P. leucopus* and assessing whether vaccination of dams with recombinant OspA-expressing *E. coli* confers protection to neonates. We tested whether vaccination of dams until breeding pairs were created, until parturition and until the pups reached 2 weeks of age influences antibody transfer and protection against tick-mediated *B. burgdorferi* challenge. Our findings have implications for reservoir-targeted vaccination strategies aimed at reducing Lyme disease risk by breaking the transmission cycle of *B. burgdorferi*.

## Materials and Methods

### Ethics Statement

This study was conducted in accordance with the Guide for the Care and Use of Laboratory Animals of the US National Institutes of Health. The protocol was approved by the University of Tennessee Health Science Center Institutional Animal Care and Use Committee (IACUC), protocol #22-0400. *Peromyscus leucopus* were sourced from the Peromyscus Genetic Stock Center (PGSC) at the University of South Carolina.

### Vaccine Preparation

Recombinant *E. coli* BL21(DE3)pLysS encoding *B. burgdorferi* OspA (*E. coli*-OspA) was produced as previously described (23). Briefly, *E. coli*-OspA was grown in Terrific Broth (TBY) at 37°C until OD600 ∼0.8, induced with 1 mM IPTG for 3 h, transferred to filter-top flasks and inactivated with 1% β-propiolactone (BPL, 98%, AlfaAesar), incubated at room temperature at 100 rpm for 24h, and harvested by centrifugation at 2000×g at 4°C. The bacterial biomass was resuspended in phosphate-buffered saline (PBS) and sprayed onto feed pellets. Pellets (RTV pellets) were allowed to dry between application layers. Inactivation was tested by inoculation of BPL-treated culture in TBY (1:200) and incubation overnight at 37°C, shaking at 250 rpm. The following day, culture inactivation was confirmed by checking turbidity and recording OD_600_=0.

### Vaccination Strategy

Dams (n=9) received RTV pellets daily for 5 consecutive days per week, with a 2-day rest period over the weekend in which they received regular mouse chow. Control dams received uncoated pellets. Dams consumed RTV pellets continuously as per vaccination schedule up to 36 weeks (**Fig 1**) and were assigned to three experimental groups based on breeding milestones. The level of IgG to OspA was analyzed in dams’ blood before selecting animals with high antibody in serum to create breeding pairs.

**Figure 1.**
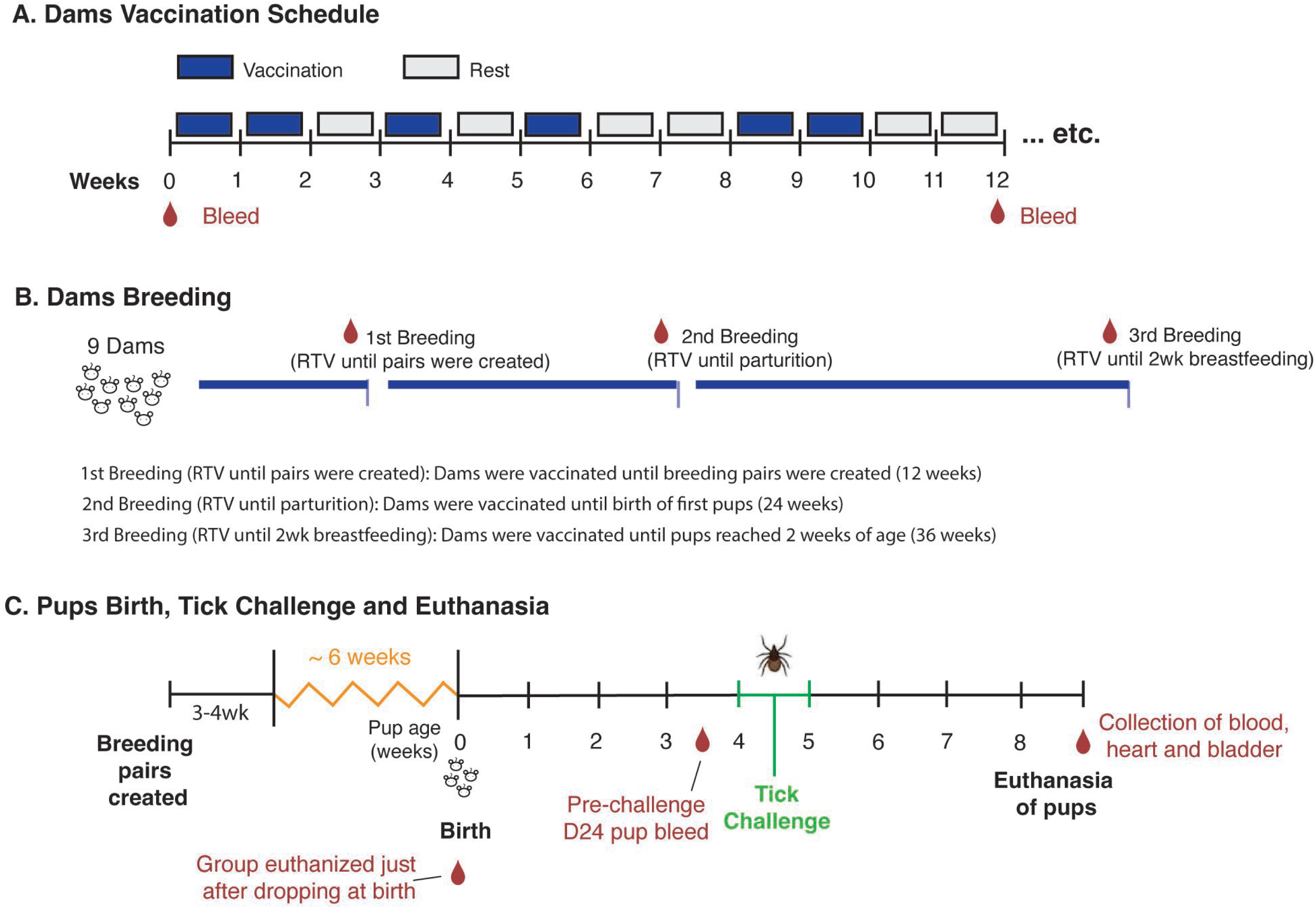
Experimental design. A) Oral vaccination schedule of adult female *Peromyscus leucopus*: mice were allowed to eat RTV pellets on vaccination weeks; pellets were exchanged for regular feed during rest weeks; control mice received uncoated pellets; B) Timeline for breeding of dams: dams were bled to evaluate serologic antibody to OspA and three experimental groups were set as 1^st^, 2^nd^ and 3^rd^ Breeding based on duration of vaccination and breeding milestones; C) Birth of pups, challenge with *B. burgdorferi* infected *I. scapularis* ticks and collection of samples.

### Breeding and Litter Management

Breeding pairs were established based on dams having anti-OspA titers (OD450>0.8) in serum diluted 1:100 assessed by ELISA. Each vaccinated female was cohoused with an unvaccinated male for 3–4 weeks, after which pregnant females were single-housed until parturition (**Fig 1**). The first group (Breeding 1) was created with 5 dams vaccinated for 12 weeks prior to breeding pairs were formed. The second group, Breeding 2, was created with 4 dams vaccinated for 24 weeks until parturition. The third group, Breeding 3, was created with the dams who maintained high levels of antibodies to OspA and remained on the vaccination regimen for 36 weeks, ensuring that immunization continued until their pups reached two weeks of age. Dams that lost the ability to reproduce or had low levels of serologic antibody to OspA were euthanized. A group of pups (n=7) born after the 3^rd^ breeding were euthanized immediately after they dropped at parturition and blood was collected. Pups remained with their dams until 24–25 days of age, at which point they underwent nymphal tick challenge.

### Blood Collection

Blood samples were collected via submandibular vein bleed. Dams were sampled at day 0 and before each breeding. Pups were sampled on the day they were born, at day 24 (pre-challenge) and day ∼71 (terminal bleed). Samples were allowed to clot for 30 min at room temperature before centrifugation at 2000×g for 10 min at 4°C. Serum was aliquoted and stored at −80°C until analysis.

### Tissue Collection and Processing

At necropsy on d71 (∼9-10 week old pups), heart and bladder tissues were harvested, transported on dry ice, and stored at −80°C. For organ culture experiments, freshly excised heart tissue was immediately placed into Barbour-Stoenner-Kelly H (BSK-H) medium supplemented with 6% rabbit serum (Sigma-Aldrich, St. Louis, MO) and incubated at 34°C for 3 weeks (on the 7^th^ day of culture, the tissue was removed to avoid putrefaction). On day 21 of culture supernatants were analyzed for motile *B. burgdorferi* detection using Petroff-Hausser chamber (dark-field microscopy, 200× magnification) and for DNA quantification via *flaB* qPCR.

### Tick Challenge and Monitoring of *B. burgdorferi* Infection

To assess susceptibility to *B. burgdorferi* infection, pups were challenged with infected nymphal *I. scapularis* ticks at ∼ 4 weeks of age. Each pup was exposed to 5–7 infected nymphs, which were allowed to feed for 72 hours in a controlled containment chamber. Flat nymphal ticks were carefully transferred from storage vials using fine forceps and placed at the base of the neck and upper back of gently restrained pups. After placement, pups were monitored to ensure proper tick attachment before being returned to their cages. After 72 hours, all engorged ticks naturally detached and were collected from the bottom of the cages. Detached ticks were stored at −20°C until further analysis for *B. burgdorferi* colonization via dark-field microscopy.

The infected nymphs used in this study were generated by feeding larval *I. scapularis* (Oklahoma Tick Laboratory) on a mouse infected with a multi-strain culture of *B. burgdorferi* sensu stricto. This *B. burgdorferi* culture was recovered from heart and bladder tissues of C3H/HeN mice infected with nymphal ticks collected in Massachusetts in 2020. To maintain infection across generations, the multi-strain *B. burgdorferi* inoculum was first introduced into C3H mice, after which successive mice were infected through tick-borne transmission. Larvae were allowed to feed on infected mice, acquire *B. burgdorferi*, and molt into infected nymphs, which were then used to challenge mice in a cycle that mimics natural enzootic transmission of *B. burgdorferi*.

### Detection of Anti-OspA and Anti-PepVF IgG by ELISA

Serum antibody responses to OspA and to *B. burgdorferi* pepVF were assessed using an enzyme-linked immunosorbent assay (ELISA). PepVF contains two peptides from *B. burgdorferi* VlsE and FlaB proteins in tandem, separated by a 3aa glycine linker, and is used in our lab to determine *B. burgdorferi* infection (24). Presence of antibody to PepVF indicates exposure to *B. burgdorferi*. Purified recombinant OspA (1 μg/mL) or recombinant PepVF (1 μg/mL) was adsorbed onto Nunc MaxiSorp plates (Thermo Fisher Scientific, Waltham, MA) by overnight incubation at 4°C. Following antigen coating, the plates were blocked with 5% bovine serum albumin (BSA) in PBS-Tween (0.05%) for 1 hour at room temperature to prevent nonspecific binding. Serum samples were diluted 1:100 in blocking buffer and incubated on the plates for 2 hours at 37°C. To detect antigen-specific IgG, a goat anti-mouse IgG antibody conjugated to horseradish peroxidase (HRP) (1:6000; Jackson ImmunoResearch, West Grove, PA) was added, followed by a 1-hour incubation at 37°C. After washing, plates were developed using TMB substrate (Sigma-Aldrich) for 15 minutes before stopping the reaction with 1N H₂SO₄. Optical density (OD) was measured at 450 nm and blanked against the control using a SpectraMax Plus ELISA reader (Molecular Devices, San Jose, CA). For Cutoff determination the positivity threshold was defined as mean + 3 SD of negative controls.

### Molecular Detection of *B. burgdorferi* DNA by qPCR

Total DNA was extracted from heart and bladder tissues and heart culture media using the DNeasy Blood & Tissue Kit (Qiagen, Germantown, MD) according to the manufacturer’s protocol. qPCR was performed to quantify *B. burgdorferi flaB* gene copies per mg of tissue or per mL of culture media. Quantitative PCR (qPCR) was performed to quantify *B. burgdorferi* flaB gene copies in DNA extracted from heart and bladder tissues, as well as culture supernatants. Reactions were carried out in a 25 μL volume using 2X TaqMan Master Mix (Thermo Fisher Scientific), with final primer and probe concentrations of 200 nM. Each reaction contained 5 μL of DNA template, and all samples were run in duplicate to ensure reproducibility. The primer and probe set used for TaqMan-based detection of the *flaB* gene was as follows: Forward primer: 5’-AAGCAATCTAGGTCTCAAGC-3’; Reverse primer: 5’-GCTTCAGCCTGGCCATAAATAG-3’; Probe: 5’-FAM-AGATGTGGTAGACCCGAAGCCGAG-TAMRA-3’. Amplification was performed using a QuantStudio 7 Real-Time PCR System (Applied Biosystems) under the following cycling conditions: an initial denaturation at 95°C for 10 minutes, followed by 45 cycles of 95°C for 15 seconds and 60°C for 1 minute. Fluorescence was measured at each cycle, and *flaB* copy numbers were determined by comparison to a standard curve generated from known concentrations of *B. burgdorferi* genomic DNA

### Statistical Analysis

Statistical analyses were conducted using GraphPad Prism 9.0 (GraphPad Software, San Diego, CA). To assess the distribution of the data, the Shapiro-Wilk test was applied to determine normality. For group comparisons, an unpaired t-test was used when data followed a normal distribution, while the Mann-Whitney U test was applied to non-normally distributed data. A significance threshold of p < 0.05 was used for all statistical tests.

## Results

### Transfer of Anti-OspA Antibodies from Dams to Pups

Pups were born to dams undergoing a continuous vaccination schedule. To determine the extent of maternal antibody transfer, serum anti-OspA IgG titers were measured in pups when they reached D24 (pre-challenge). Pups born from dams vaccinated until breeding pairs were formed are grouped under 1^st^ breeding. Pups born from dams vaccinated until parturition are grouped under 2^nd^ breeding. Pups born from dams vaccinated until the pups were two weeks old are grouped under 3^rd^ breeding. The results are shown in **Fig 2**. Anti-OspA IgG was significantly increased in D24 pups born from vaccinated dams during the 1^st^, the 2^nd^ and the 3^rd^ breeding, in contrast to pups born to control unvaccinated dams. Nevertheless, we note that pups born from the 1st breeding of vaccinated dams had an average anti-OspA IgG OD_450_∼ 0.271 (SD± 0.094); pups born from the 2nd breeding of vaccinated dams had an average anti-OspA IgG OD_450_∼0.809 (SD± 0.073); and pups born from the 3rd breeding of vaccinated dams had an average anti-OspA IgG OD_450_∼1.532 (SD± 0.591). The data shows that continued vaccination of the dams past parturition maintained sufficient levels of protective antibody (OD450>0.8) in blood.

**Figure 2.**
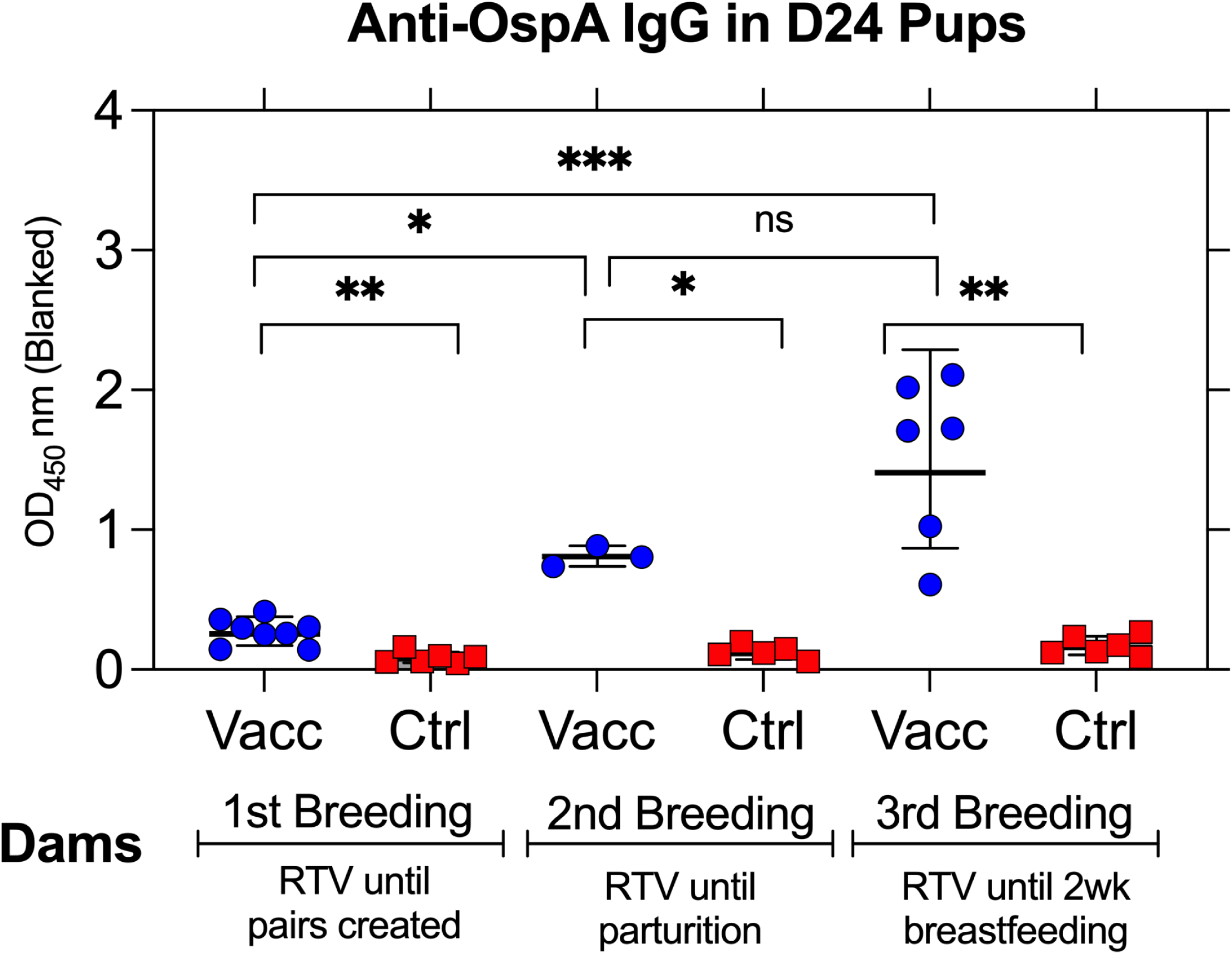
Transfer of OspA antibodies from dams to pups. ELISA quantification of pre-challenge anti-OspA IgG in sera from D24 pups born to vaccinated dams. Optical density (OD) at 450nm (blanked) is shown for each group. Each point represents an individual pup, with lines indicating group means ± SD. Statistical analysis by Mann-Whitney test *p<0.05, ** p<0.005, *** p<0.0005.

### Seroconversion to *B. burgdorferi* in Tick Challenged Pups

To assess whether pups were protected or infected with *B. burgdorferi* after tick challenge, anti-PepVF IgG was measured in serum samples collected at euthanasia (∼day 71), at least 3 weeks after the last day of challenge (**Fig 3).** Pups (n=8) from the 1^st^ Breeding of vaccinated dams had equivalent anti-PepVF IgG as control pups (n=10), confirming infection with tick-transmitted *B. burgdorferi* in both groups (p = 0.7642) (**Fig 3A**). Although 2/3 pups from the 2^nd^ Breeding of vaccinated dams did not have anti-PepVF antibodies (**Fig 3B**, circle), overall differences with the control group (n=5) were not significant (p = 0.9432). Strikingly, pups (n=6) from the 3^rd^ Breeding of vaccinated dams (**Fig 3C**) had anti-PepVF IgG levels below the cutoff in contrast to the control group (n=6), p = 0.0147, indicating absence of infection. These results suggest that maternal vaccination that extended until and beyond parturition conferred anti-OspA passive immunity to pups.

**Figure 3.**
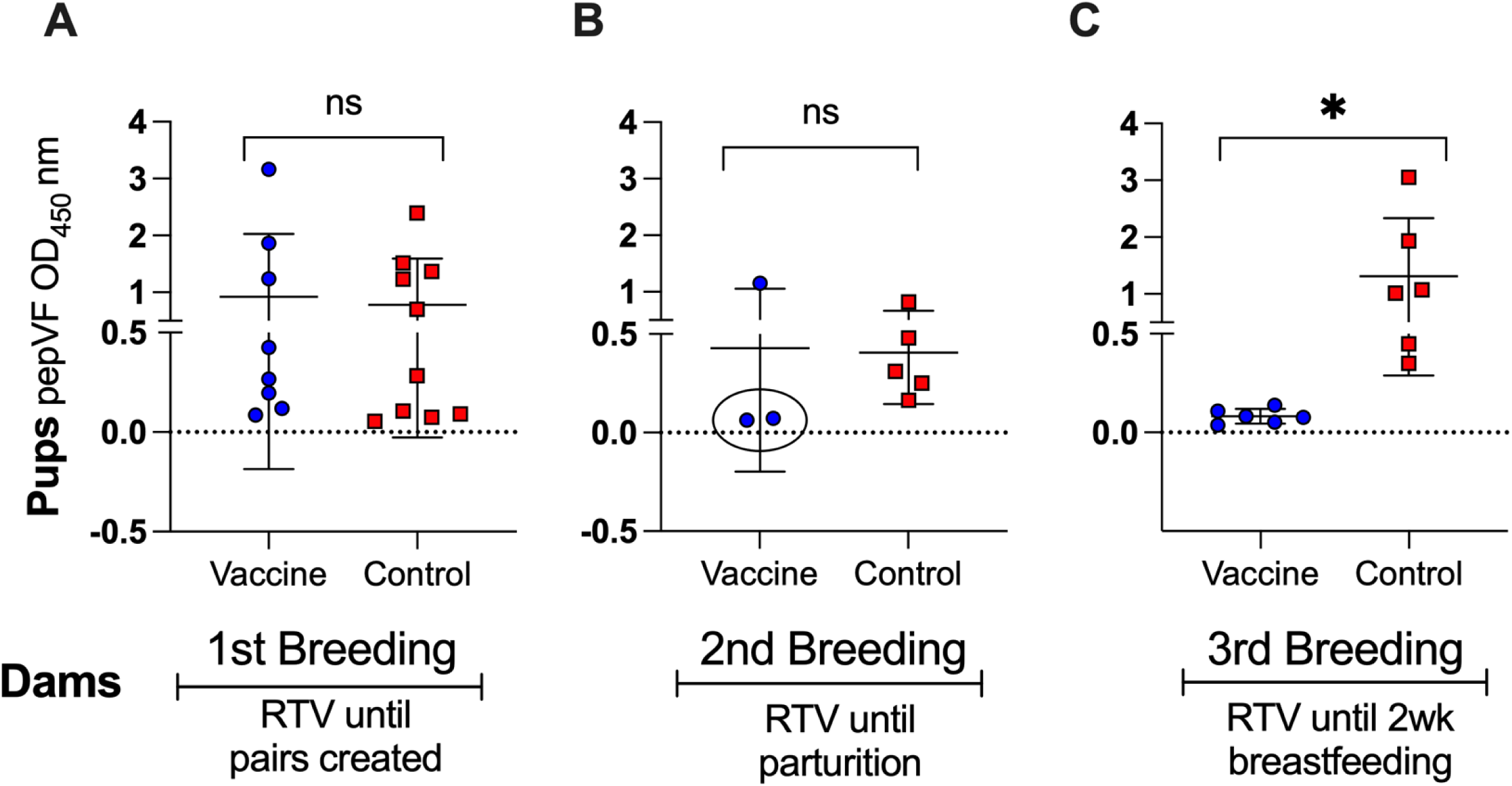
Seroconversion to *B. burgdorferi* in tick challenged pups. (A–C) ELISA quantification of anti-PepVF IgG in terminal sera from pups born to vaccinated dams. Pups were bled ∼ 4 weeks post tick challenge. Optical density (OD) at 450 nm (blanked) is shown for each group. Each data point represents an individual pup, with lines indicating group means ± SD. Statistics by Unpaired t test: ns, not significant, * p<0.05.

### Burden of *B. burgdorferi* in Tissues from Tick Challenged Pups

To provide additional evidence of *B. burgdorferi* dissemination, we purified DNA from heart and bladder organs from pups euthanized > 4 weeks after tick challenge (∼ day 71) and performed qPCR targeting the *B. burgdorferi flaB* gene (**Fig 4**). In pups from the 1^st^ Breeding, there are no differences in *flaB* loads in heart tissues from vaccinated and control groups; in pups from the 2^nd^ Breeding, the same 2 mice (2/3) in the vaccinated group that did not seroconvert to *B. burgdorferi*, also did not have *flaB* in heart tissues, whereas 5/5 mice in the control group had *flaB* DNA in heart tissues. Differences between the groups are not significant (p=0.3393). However, in pups from the 3^rd^ Breeding, no detectable *flaB* copies were found in bladder tissues from the vaccinated group (0/6), in contrast to the pups from the control group that were all positive (6/6) for *B. burgdorferi flaB* (p=0.0022). Absence of *B. burgdorferi flaB* DNA in pups from the 2^nd^ and 3^rd^ Breeding correlate with absence of PepVF seroconversion, further confirming the protective effect of maternal antibody transfer when dams were vaccinated until and beyond parturition.

**Figure 4.**
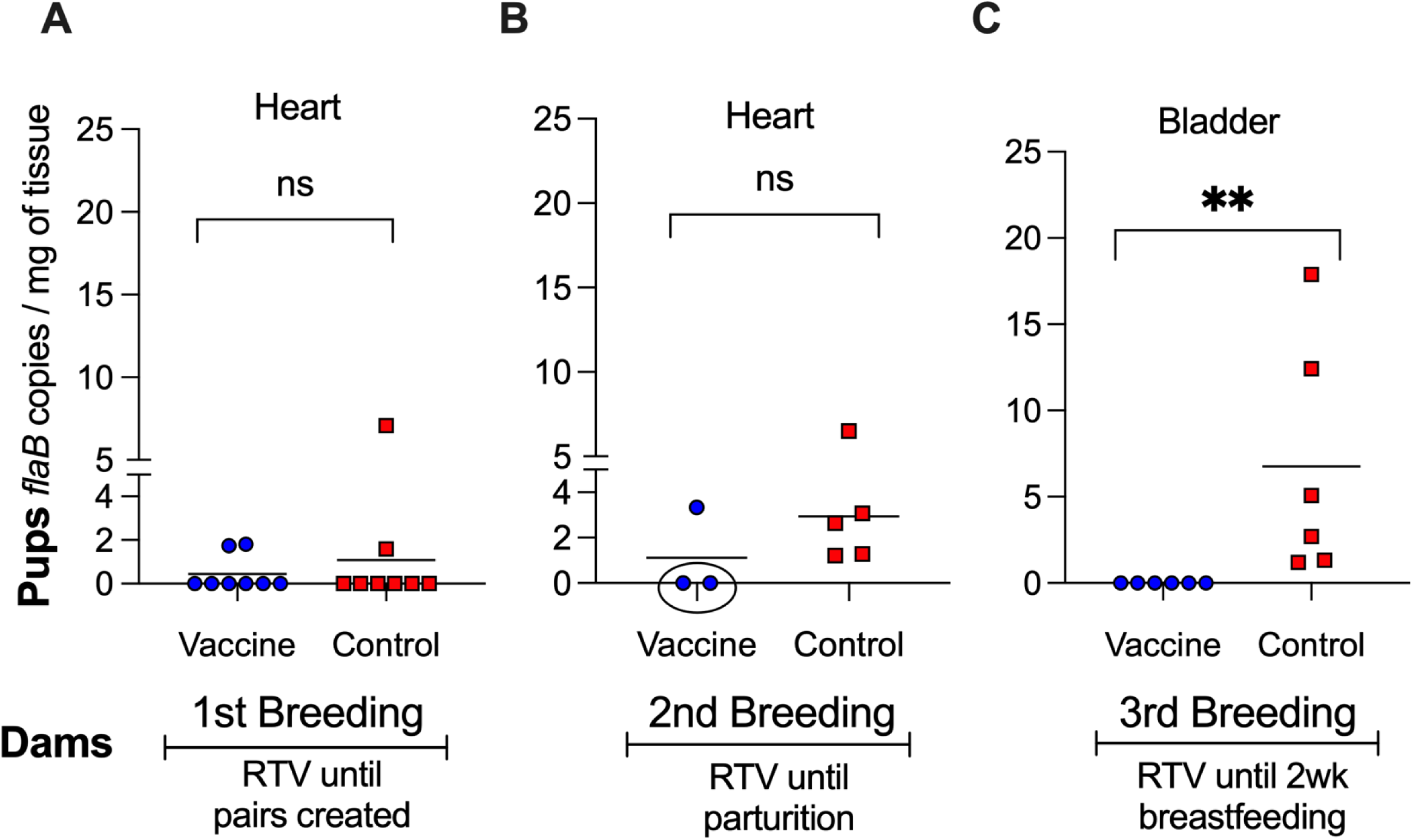
Burden of *B. burgdorferi* in tissues from tick challenged pups. (A–C) qPCR quantification of the *flaB* gene in heart and bladder tissues collected from pups ∼ 4 weeks post tick challenge. Each data point represents an individual pup, with lines indicating group means. Statistics by Mann Whitney test: ns, not significant, **p<0.005.

### Viability of *B. burgdorferi* in Tissues from Tick Challenged Pups

To further assess *B. burgdorferi* persistence, heart tissues from pups born from 3^rd^ Breeding dams were cultured in BSK-H medium at 34°C for 21 days, and live spirochetes were counted using a Petroff-Hausser chamber under a dark field microscope (**Figure 5A**). Only pups (4/5) from unvaccinated control dams had detectable motile *B. burgdorferi* in culture, confirming that live spirochetes had successfully colonized cardiac tissue after tick challenge. The control group exhibited a range of 10^5^-10^6^ spirochetes per mL of culture, while pups from vaccinated dams had no visible live bacteria, p=0.0152. qPCR analysis of heart culture supernatants (**Figure 5B**) confirmed that high *flaB* copy numbers (mean 10^5^) were detected in control heart cultures (5/5), whereas no *B. burgdorferi* DNA was present in pups from vaccinated dams (0/5), p=0.0022. This indicates that dams that received vaccine until 2 weeks post-parturition, produced pups that were protected from cardiac colonization by *B. burgdorferi*.

**Figure 5.**
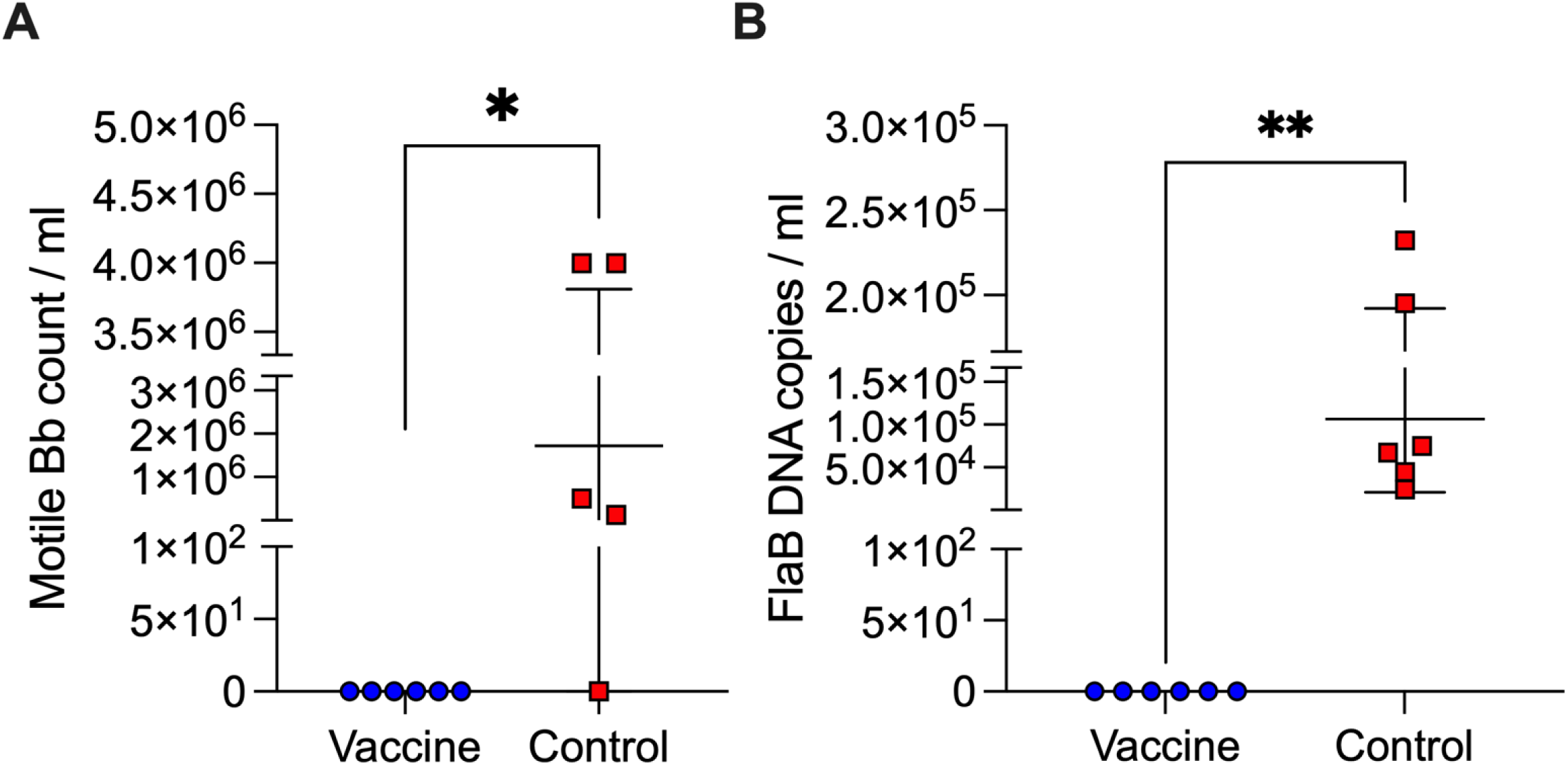
Viability of *B. burgdorferi* in tissues from tick challenged pups. Quantification of *B. burgdorferi* in cultures of heart tissue from pups at ∼ 4 weeks post-challenge: (A) Enumeration of motile *B. burgdorferi* using a Petroff-Hausser chamber under a dark field microscope; (B) qPCR quantification of *B. burgdorferi flaB* in heart culture media. Each data point represents an individual pup, with lines indicating group means ± SD. Statistics by Mann Whitney test, *p<0.05 and ** p<0.005.

### Absence of OspA IgG in Serum from Pups Euthanized After Birth

To determine whether maternal antibodies were transferred in utero, OspA ELISA was performed on serum from pups euthanized immediately after they dropped at birth in comparison to the respective dam (**Fig 6**). In contrast to the respective dam, no anti-OspA IgG was detected in newborns from the vaccinated dam, confirming that placental transfer of antibodies was absent.

**Figure 6.**
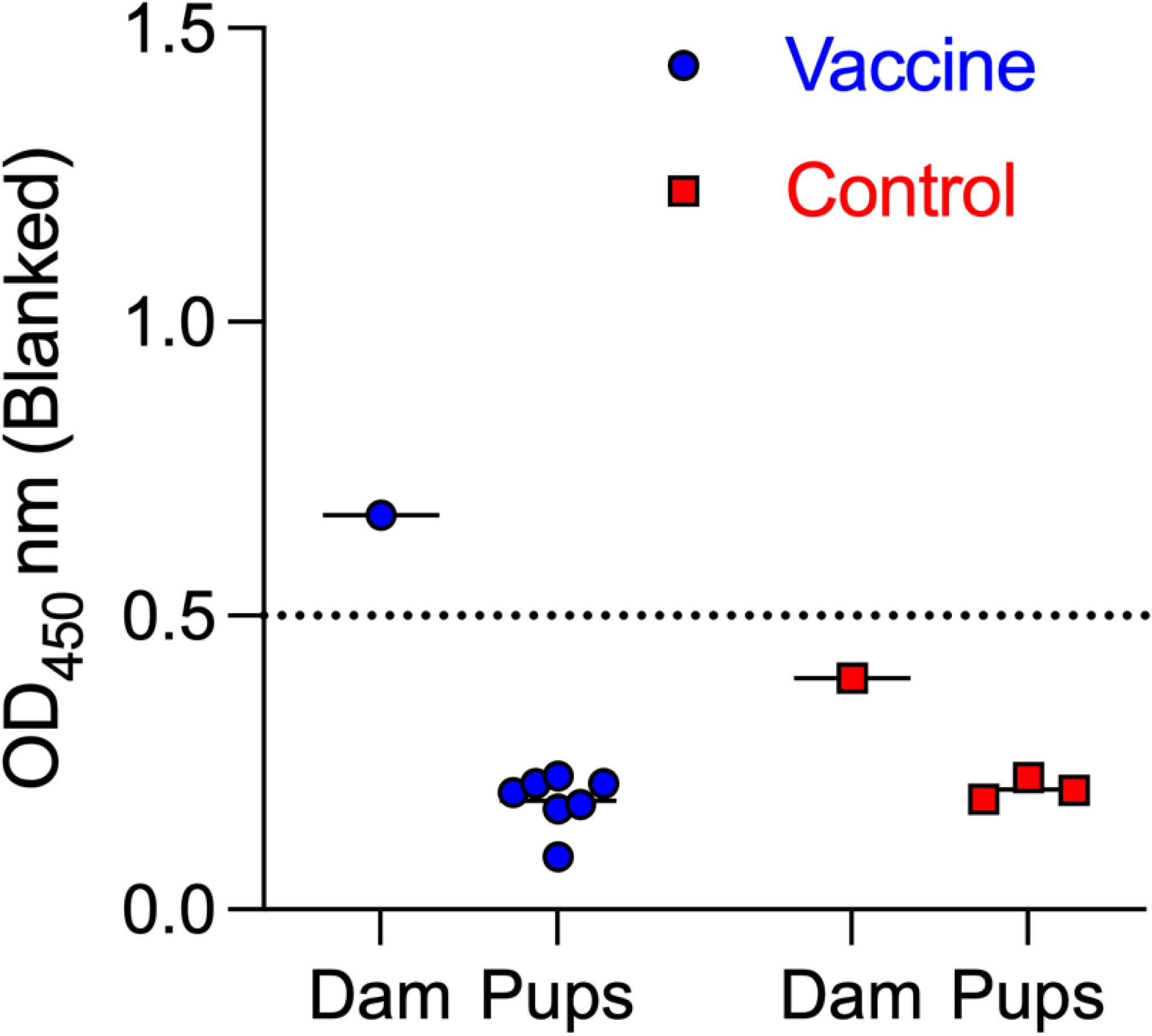
Absence of OspA IgG in serum from pups euthanized after birth. ELISA quantification of anti-OspA IgG in serum from pups euthanized immediately after birth from dams vaccinated until two weeks parturition (3^rd^ Breeding). Optical density (OD) at 450 nm (blanked) is shown. Each data point represents an individual pup, with lines indicating group means. The cutoff value (dashed line) represents the mean + 3 SD of negative controls.

### Anti-OspA IgG depletion from vaccinated dam serum after lactation

To investigate whether maternal antibody depletion occurred due to antibody transfer to offspring, serum anti-OspA IgG were measured in dams used for the 3^rd^ breeding at two time points (11Jun and 14Oct) and in the respective litters, on the latter timepoint, before tick challenge (**Fig 7**). We found that the reduction in OspA antibody measured between the two time points in the dams corresponds to the increase in OspA antibody measured in the respective litters in the latter timepoint: ∼0.7OD_450_ for Dam 1 (**Fig 7A**) and ∼1.8OD_450_ for Dam 2 (**Fig 7B**). These findings suggest that transfer of antibody from dam to pups may be dependent on the efficiency of the pups’ suckling.

**Figure 7.**
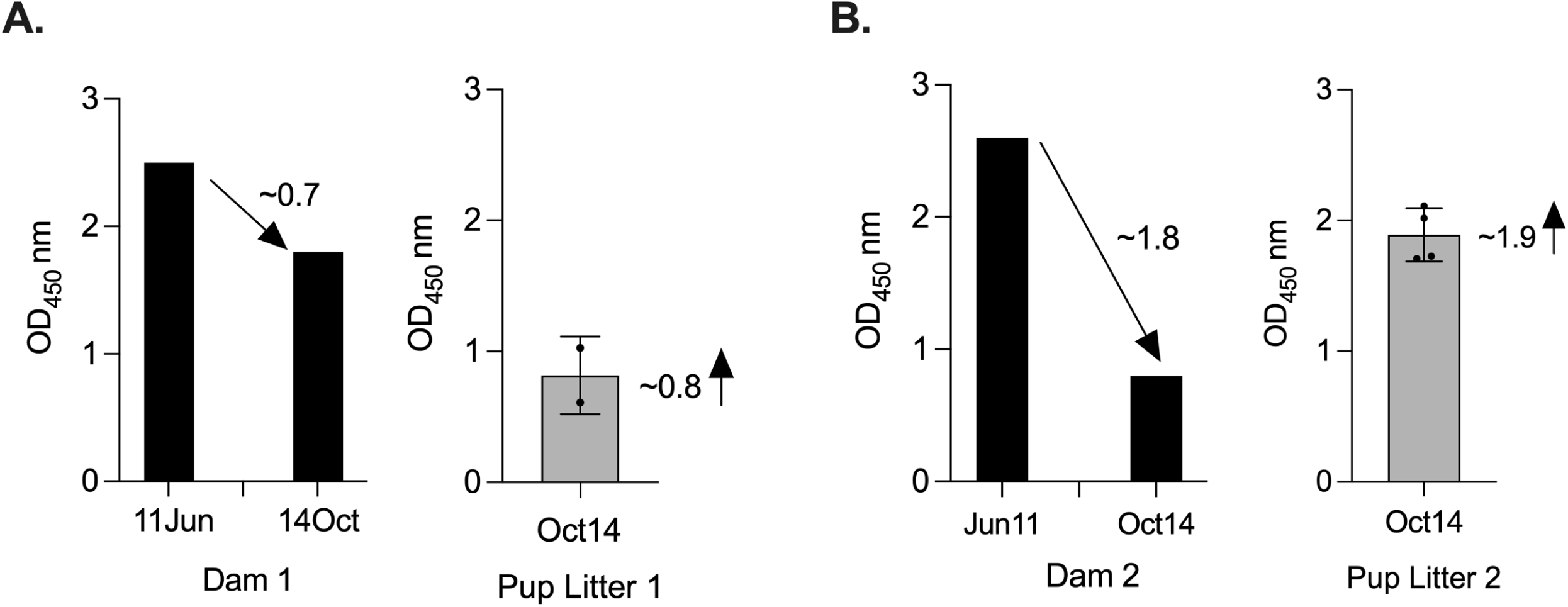
Anti-OspA IgG depletion from vaccinated dam serum after pup suckling. ELISA quantification of anti-OspA IgG in serum from two dams used in the 3rd breeding and the respective litters. Sera from dams were collected at two time-points ∼ 4 months apart. Sera from the respective litters were collected on the latter time point, before tick challenge. Each data point represents an individual pup, with lines indicating group means ± SD.

## Discussion

Our study demonstrates that vaccination of *Peromyscus leucopus* dams against *B. burgdorferi* lead to transfer of anti-OspA IgG to offspring and that, the extent of transfer and protective efficacy depends on maintenance of maternal immunization. Pups born to dams vaccinated until parturition and until two weeks post-parturition exhibited significantly higher anti-OspA IgG levels than those from dams vaccinated until breeding pairs were created. After challenge with *B. burgdorferi* infected *I. scapularis* ticks, pups born to vaccinated dams that maintained high systemic anti-OspA IgG had no anti-*B. burgdorferi* antibody in serum, no *B. burgdorferi* DNA in bladder or heart tissues and no motile *B. burgdorferi* in heart tissue. Furthermore, we detected no serologic anti-OspA IgG in offspring from vaccinated dams, euthanized immediately after birth, before suckling. The data indicates that anti-OspA antibody is transferred from *P. leucopus* dams to pups via lactation and that it protects against tick-transmitted *B. burgdorferi*.

Vaccination of *P. leucopus* dams until the breeding pairs were created (1^st^ Breeding) did not protect *P. leucopus* pups from tick-mediated infection due to absence of sufficient levels of anti-OspA antibodies (OD_450_<0.8, **Fig 2**) in pups born ∼ 10 weeks after the dams were vaccinated. This was surprising given our previous studies in *P. leucopus* (25) and in C3H/HeN mice (22) in which we vaccinated mice with the reservoir targeted vaccine (RTV). In the first study, we vaccinated *P. leucopus* with 20-30 units of live RTV for 1-4 months and found that mice maintained anti-OspA antibody for up to a year (25). In the second study, we found that C3H/HeN dams vaccinated with 20 units of live RTV over a period of 8 weeks, produced pups 33-79 days after vaccination that maintained protective titers of OspA antibody from 2-9 weeks after birth (22). The major difference between the studies is that in the current study we used RTV inactivated with BPL, whereas in the former studies we used live RTV. This change was intentional to evaluate a vaccine that mimics the product currently licensed by USDA, which is inactivated with BPL (26). This data indicates that production of anti-OspA antibody by *P. leucopus* requires persistent vaccination, if the vaccine formulation is inactivated likely because it does not colonize the mouse gut.

Previous studies show that transfer of maternal IgG provides passive immunity but does not necessarily confer sterilizing protection against vector-borne pathogens (20, 21). The degree of IgG transfer and its impact on early-life immunity can vary depending on maternal antibody titers, the mechanism of transfer (transplacental vs. lactogenic), the duration of exposure (27) and neutralizing capability of the antibody (25). In our study, the neutralizing capability of anti-OspA antibody transferred from *P. leucopus* dams to its pups is evidenced by the absence of *B. burgdorferi* antibody in serum of pups (2/3 in Breeding 2 and 6/6 in Breeding 3, **Fig 3**), as well as absence of DNA in heart and bladder tissues (the same mice 2/3 in Breeding 2, and 6/6 in Breeding 3, **Fig 4**) and complete absence of motile spirochetes in cultures from heart tissue (6/6 in Breeding 3, **Fig 5**). All pups from breeding 1, born to vaccinated and control dams, had antibodies to *B. burgdorferi* antigens after tick challenge (**Fig 3**) but we did not detect *B. burgdorferi* DNA in heart tissue of these mice (**Fig 4**). We perform two independent tests (antibody to *B. burgdorferi* in serum and *B. burgdorferi* DNA in tissues) and we accept one positive result as evidence of *B. burgdorferi* dissemination (24). This method offsets risks incurring from tracking all datapoints in experiments that can last up to 1 year. Thus, for Breeding 1, we consider all mice in this experimental group to be infected and that this was associated with insufficient production of anti-OspA antibody by the respective vaccinated dams. Final efficacy of our intervention is determined by culture of *B. burgdorferi* from tissues of tick-challenged mice. **Fig 5** shows conclusively that pups born to dams vaccinated until 2 weeks post-parturition did not have viable *B. burgdorferi* in cultures from heart tissue. This shows that for tick-transmitted *B. burgdorferi,* if *P. leucopus* is persistently vaccinated and produces sufficient anti-OspA antibody to be transferred to pups, the offspring is protected to at least 4 weeks post parturition. In addition, we show that immediately after birth, pups do not have anti-OspA antibodies in serum (**Fig 6**). Thus, we infer that *P. leucopus* pups acquire OspA antibodies during suckling. This observation aligns with our previous studies in C3H mice demonstrating that anti-OspA IgG transfer via milk is a primary mechanism of passive immunity (22).

A potential question may arise from differences in **Fig 2** data showing significant increase of anti-OspA IgG in pups euthanized on the day of birth from dams vaccinated until parturition in Breeding 2 and data in **Fig 6** showing absence of anti-OspA IgG in pups euthanized immediately after birth in Breeding 3. This is explained by the fact that pups from the 2^nd^ breeding were born overnight and were euthanized in the morning. Thus, for pups born from the 3^rd^ breeding and to confirm if antibody was transferred via lactation, as we showed previously for C3H mice (22), we euthanized the pups immediately after they dropped to prevent any potential suckling time.

The decline in maternal anti-OspA IgG over time, particularly during lactation (**Fig 7**), suggests a depletion of circulating IgG that could have impacted the duration of passive protection provided to pups. This observation is consistent with findings in other models where maternal antibody levels decrease postnatally as antibodies are transferred to offspring (20).

From a reservoir-targeted vaccine intervention perspective, our findings suggest that if maternal anti-OspA antibodies reduce tick colonization efficiency or delay early-stage infection in *P. leucopus*, this may have broader ecological implications by disrupting pathogen transmission cycles. Further studies are needed to assess the effect of passively transferred anti-OspA antibody in reduction of nymphal infection prevalence.

## Acknowledgments

Research reported in this publication was supported by the National Institute of Allergy and Infectious Diseases of the National Institutes of Health under Awards Number R01AI139267 (MGS), R43AI155211 (MGS) and R44AI167605 (MGS). The content is solely the responsibility of the authors and does not necessarily represent the official views of the National Institutes of Health.

License: CC-BY-ND

## Notes

### Competing Interest Statement

MGS has competing interests that relate to a licensed patent. The other authors have no conflicts.

### Summary of Updates

The text and the figures have been revised according to the reviewers recommendations. No supplemental material is included.

